# Growth-factor like gene regulation is separable from survival and maturation in antibody secreting cells

**DOI:** 10.1101/439752

**Authors:** Sophie Stephenson, Matthew A Care, Im Fan, Alexandre Zougman, David R Westhead, Gina M Doody, Reuben M Tooze

## Abstract

Recurrent mutational activation of the MAP kinase pathway in plasma cell myeloma implicates growth factor-like signaling responses in the biology of antibody secreting cells (ASCs). Physiological ASCs survive in niche microenvironments, but how niche signals are propagated and integrated is poorly understood. Here we dissect such a response in human ASCs using an in vitro model. Applying time course expression data and parsimonious gene correlation networking analysis (PGCNA), we map expression changes that occur during the maturation of proliferating plasmablast to quiescent plasma cell under survival conditions including the potential niche signal TGFB3. This analysis demonstrates a convergent pattern of differentiation, linking UPR/ER stress to secretory optimization, co-ordinated with cell cycle exit. TGFB3 supports ASC survival while having a limited effect on gene expression including up-regulation of CXCR4. This is associated with a significant shift in response to SDF1 in ASCs with amplified ERK1/2 activation, growth factor-like immediate early gene regulation and EGR1 protein expression. Similarly, ASCs responding to survival conditions initially induce partially overlapping sets of immediate early genes, without sustaining the response. Thus, in human ASCs growth factor-like gene regulation is transiently imposed by niche signals but is not sustained during subsequent survival and maturation.

## Introduction

The generation and maintenance of functional antibody secreting cells (ASCs) is essential to humoral immunity (1). Long-lived ASCs persist as plasma cells (PCs) in a range of different niche conditions in vivo, affording the potential to link sustained survival to phenotypic and functional diversity (2, 3). In PC neoplasia, abnormalities in the niche play a key role in sustaining the neoplastic clone (4), while targets of recurrent mutation in PC neoplasia identify pathways of potential functional importance to the biology of ASCs. One of the most frequently deregulated pathways is that of RAS/RAF/MAP kinase signaling (5).

Although it is widely accepted that PCs may exist in complex microenvironments across the spectrum of normal to neoplastic states, how the pattern of signals received by a PC may be integrated remains poorly understood. We and others have developed model systems allowing the generation and maintenance of long-lived human PCs in vitro, which provide tools to directly address this question in primary cells, and link external cues to specific response pathways (6, 7). PCs are functionally defined as ASCs which have entered cell cycle quiescence and derive from a preceding proliferative ASC state referred to as plasmablasts (PBs). This transition is accompanied by phenotypic changes but is principally separated by entry into cell cycle quiescence (8). The ability of an ASC to survive as a PC, can be conceptually reduced to the capacity of the cell to home to, reside in and respond to relevant niche signals, and it has been argued that competition for niche residence may contribute to control of the long-lived PC pool (9, 10). The chemokine CXCL12/SDF1 has been identified as an important component of niche homing signals for PCs (11–13). Consequently SDF1-rich mesenchymal stromal cells are considered to form an important element of the marrow niche (14, 15). In addition to secreting SDF1, bone marrow stromal cells have the capacity to secrete a diverse range of mediators amongst which is TGFB (4). Cross-talk between TGFB and SDF1 signaling pathways has been described in several cell systems (16, 17). Both pathways are involved in the process of epithelial-mesenchymal transition, and hence with invasive and migratory behavior (18, 19). However, whether PCs integrate these signals and to what effect is not known.

Amongst the signaling pathways linked to SDF1 responses in lymphocytes is activation of the MAP kinase pathway (16, 20–22). Although the role of the MAP kinase pathway in normal PC biology is not defined, components of the pathway are recurrent targets of mutation in PC neoplasia including both upstream regulators such as the RAS oncogenes and downstream effector EGR1 (23–25). EGR1 mutation has been reported to show a particularly high cancer clonal fraction when mutated, suggesting that it may either exert a strong selective pressure or be an early event in pathogenesis (5). Interestingly in a model of cell cycle progression established in human mammary epithelial cells, ERK-EGR1 signaling has been proposed to provide a threshold mechanism generating all-or-none decisions for cell cycle entry (26). Furthermore, EGR1 protein expression along with several other immediate early genes can act as a sensor for the duration of MAP kinase signaling (27–29).

Here we analyze TGFB3 and SDF1 responses in human PCs using time course expression data and network analysis. This provides evidence for a model of convergent differentiation largely independent of the conditions supporting PC survival during the transition from PB to quiescent PC state. SDF1 exposure provides a pulse of MAP kinase signaling, which can be significantly enhanced in the presence of TGFB3. This translates into enhanced induction of immediate early genes and EGR1 protein expression, mimicking classical growth factor responses. ASCs responding to in vitro survival conditions induce transient and partially overlapping sets of immediate early genes, but without sustaining the response. These data indicate that ASCs responding to niche signals follow rules for growth factor-induced MAP kinase signaling established in model epithelial and neuronal cell lines, and that growth factor-like gene regulatory responses are a transient feature of the ASC response to niche signals.

## Materials & Methods

### Reagents

IL2 (Miltenyi); IL21, IL6, IFNα, TGFB3, SDF1α, Osteoprotegerin, M-CSF, Osteopontin, SCF, MIF, IGF-BP2, IGF-II, and Activin A (Peprotech); Jagged-1, ANGPT1, VEGF-165 and VEGF-121 (R&D); Multimeric-APRIL (Caltag); Goat anti-human IgM & IgG F(ab’)_2_ fragments (Jackson ImmunoResearch); Hybridomax hybridoma growth supplement (Gentaur); Lipid Mixture 1, chemically defined (200X) and MEM Amino Acids Solution (50X) (Sigma); SB525334 (Selleckchem).

### Donors and cell isolation

Peripheral blood was obtained from healthy donors after informed consent. The number of donors per experiment is indicated in the figure legend. Mononuclear cells were isolated by Lymphoprep (Axis Shield) density gradient centrifugation. Total B-cells were isolated by negative selection with the Memory B cell Isolation Kit (Miltenyi).

Bone marrow aspirates were from 3 anonymous donors and were derived from surplus clinical samples. Mononuclear cells were isolated by Lymphoprep and normal B cells and plasma cells identified by the gating strategy outlined in Supplementary Figure SF1A.

### Cell cultures

Cells were maintained in 24-well flat-bottom culture plates (Corning) and Iscove’s Modified Dulbecco Medium (IMDM) supplemented with Glutamax and 10% heat-inactivated fetal bovine serum (HIFBS, Invitrogen), Hybridomax hybridoma growth supplement (11 μl/ml), Lipid Mixture 1, chemically defined and MEM Amino Acids solution (both at 1x final concentration) from day 3 onwards.

Day 0 to day 3 - B cells were cultured at 2.5 × 10^5^/ml with IL2 (20 U/ml), IL21 (50 ng/ml), F(ab’)_2_ goat anti-human IgM & IgG (10 μg/ml) on γ-irradiated CD40L expressing L-cells (6.25 × 10^4^/well).

Day 3 to day 6 - At day 3, cells were detached from the CD40L L-cell layer and reseeded at 1 × 10^5^/ml in media supplemented with IL2 (20 U/ml) and IL21 (50 ng/ml).

Day 6 onwards - For cytokine combination experiments and TGFB3 dose response, cells were seeded at 5 × 10^5^/ml in media supplemented with IL6 (10 ng/ml), IL21 (50 ng/ml), IFNα (100 U/ml) and combinations of Jagged 1 (2 μg/ml); Osteopontin (1 μg/ml); Osteoprotegerin, SCF and IGF-BP2 (100 ng/ml); Activin A, and M-CSF (50 ng/ml); IGF-II (30 ng/ml): ANGPT1, VEGF-165, VEGF-121, MIF and TGFB3 (10 ng/ml or 0.1-1000 ng/ml for dose response). Cell culture was terminated at day 13. For inhibitor experiments, cells were seeded at 5 × 10^5^/ml in phenol red free, serum free IMDM (Invitrogen) for 18h. Vehicle or SB525334 (1 μM) was added for 1h and then cells were stimulated with either IL6 (10 ng/ml), IL21 (50 ng/ml) and IFNα (100 U/ml) or TGFB3 (2.5 ng/ml) for 2h.

For extended gene expression experiments, cells were reseeded at 1 × 10^6^/ml in media supplemented with IL6 (10 ng/ml), IL21 (50 ng/ml) and either IFNα (100 U/ml) or TGFB3 (2.5 ng/ml) alone or in combination. Cells were refed at 3.5-day intervals and analyzed at indicated times.

For short gene expression experiments, cells were seeded at 1 × 10^6^/ml in phenol red free media supplemented with 0.5% HIFBS, IL6 (10 ng/ml), IL21 (10 ng/ml) alone or with TGFB3 (2.5 ng/ml) for 20 hours. SDF1 (1 ng/ml) was added and cells analyzed at indicated times.

### Flow cytometric analysis and microscopy

Cells were analyzed using 4- to 6-color direct immunofluorescence staining on a BD LSR II (BD Biosciences) or Cytoflex S (Beckman Coulter) flow cytometer. Antibodies used were: CD19 PE (LT19), CD19 VioBlue (LT19), CD138 APC (B-B4/44F9) and CD138 VioGreen (44F9) (Miltenyi); CD20 e450 (2H7) (eBioscience); CD27 FITC (M-T271), CD56 PECy7 (NCAM16.2), CD38 PECy7 (HB7), CD38 BUV395 (HB7) and phospho-Smad2/3 PE (BD Biosciences); CXCR4 PE (R&D Systems). Phosflow Lyse/Fix and Perm/Wash buffer (BD Biosciences) was used for the preparation of cells prior to Phosflow. Controls were isotype-matched antibodies or FMOs. Dead cells were excluded by 7-AAD (BD Biosciences). Absolute cell counts were performed with CountBright beads (Invitrogen). Cell populations were gated on FSC and SSC profiles for viable cells determined independently in preliminary and parallel experiments. Analysis was performed with BD FACSDiva Software 5.0 and Flowjo version 10 (BD Biosciences).

### RNA, cDNA and RT-PCR

RNA was extracted with TRIzol (Invitrogen), subjected to DNAseI treatment (DNAfree, Ambion) and reverse transcribed using Superscript II Reverse Transcriptase (Invitrogen). Taqman^®^ Assays for *FOS* (Hs00170630_m1), *FOSB* (HsOO 17185l_m1), *EGR1* (Hs00152928_m1) and *PPP6C* (Hs00254827_m1) were carried out according to manufacturer’s instructions and run on a Stratagene Mx3005p.

### Protein analysis

At the indicated time points, cells were lysed in Laemmli buffer. Samples were separated by SDS-PAGE and transferred to nitrocellulose. Proteins were detected by ECL (SuperSignal WestPico PLUS or Fento Chemiluminescent substrate, Thermo Scienctific) and visualized on a ChemiDoc (BioRad) or film. Protein bands were quantitated using ImageJ software.

Antibodies used were β-ACTIN (Sigma); p-SMAD2 S465/467 (138D4), p-ERK1/2, ERK1/2 and EGR1 (Cell Signaling); goat anti-mouse HRP, goat anti-rabbit HRP (Jackson ImmunoResearch).

### Proteomics

Centrifuged and filtered (0.2 μm) supernatants and media were generated from duplicate samples of M2-10B4 cells either non-irradiated, irradiated or irradiated and cultured in the presence of IL6, IL21 and IFN⍰. STrap-based tryptic digestion was performed as previously described (30). Peptides were separated online by reversed-phase capillary liquid chromatography using EASY-nLC 1000 UPLC system (Thermo) connected to a capillary emitter column. LTQ-Orbitrap Velos mass spectrometer (Thermo) was used for data acquisition. Raw data were processed against the Uniprot mouse protein sequence database (May, 2013) using MaxQuant software (www.maxquant.org) (31). The maximum protein and peptide false discovery rates were set to 0.01.

### Immunohistochemistry

Surplus material from bone marrow trephine specimens assessed as reactive in diagnostic practice were stained with phospho-SMAD2 specific antibody (Cell Signalling rabbit polyclonal, cat no. 3101). Staining was performed using DAKO Envision Flex and Detection System (Envision Flex+ high pH kit (K8002 DAKO)). Antigen retrieval was carried out using heat mediated antigen retrieval using pressure cooking for 6 min. DAKO Envision Flex+ kit was used with standard methods, Flex+ reagent with rabbit linker, DAB chromogen and haematoxylin counter-stain (complete protocol available on request). Stained slides were assessed on a Nikon Eclipse 80i microscope equipped with x40 Plan Fluor objective and Nikon Digital Sight DS-Fi1 camera system.

### Expression data sets

Two gene expression data sets were generated, a long time course (LTC) and a short time course (STC). The LTC consists of a differentiation of B cells from 4 healthy donors from day 3 (activated B-cell), day 6 (PB) and then post treatment with three different conditions at day 6 +1h, +3h, +6h, +12h, +24h (day 7), +48h (day 8), +96h (day 10), +168h (day 13; PC), +336h (day 20; PC). The three conditions were, 1: (IL6, IL21, IFNα), 2: (IL6, IL21, IFNα, TGFB3) and 3: (IL6, IL21, TGFB3.). For each time there were 4 samples except for day 6 (biological replicates giving 8 samples) and condition 3 day 20 which only had 1 sample due to quality control. The STC consists of differentiating PBs (day 7) from 3 healthy donors maintained in low serum media with/without TGFB and then post treatment with SDF-1 at +30min, +120min, +360min.

### Gene expression data acquisition and analysis

Gene expression analysis was performed using HumanHT-12 v4 Expression BeadChips (Illumina) according to the manufacturer’s instructions and scanned with the Illumina BeadScanner. Initial data analysis was performed using GenomeStudio Gene Expression Module, and as previously described. (32)

### Gene signature data and enrichment analysis

A merged data set of 39,482 terms was generated from externally curated data bases and gene ontology terms as previously described. (33) Enrichment of gene lists for signatures was assessed using a hypergeometric test, in which the draw is the gene list genes, the successes are the signature genes, and the population is the genes present on the platform.

### PGCNA network analysis

#### Probe selection

Before probes were merged per gene the most informative probes were selected for the different time courses. For the LTC the probes differentially expressed between each condition at each time point and between day 3 and day 6 (*p*-value < 0.01; Fold-change > 1.2; n=7,073 probes) were merged with the probes σ^2^ > 0.05 (across median values per time point) giving a non-redundant set of 11,388 probes, merged to 9,063 genes. For the STC the probes differentially expressed between TGFB+/− at each time point and between each post-treatment time point and its corresponding pre-treatment sample (*p*-value < 0.05; Fold-change > 1.2; n=4,471 probes) were merged with the probes σ^2^ > 0.05 (across median values per time point) giving a non-redundant set of 5,302 probes, merged to 4,705 genes.

#### Network analysis

For details and validation of the Parsimonious Gene Correlation Network Analysis (PGCNA) approach see our other work (33). Here a brief description of the method will be give (see Figure 3A for overview). After informative genes were selected they were used to calculate Spearman’s rank correlations for all gene pairs using the Python scipy.stats package. For each gene (row) in a correlation matrix only the 3 most correlated edges per gene were retained. The resulting matrix *M*, with entries written as *M* = (*m_ij_*) was made symmetrical by setting *m_ij_* = *m_ji_* for all indices *i* and *j* so that *M* = M^T^ (its transpose). This reduced the edges in LTC from 41,064,453 to 24,683 and in STC from 11,066,160 to 11,458. The correlation matrices were clustered 10,000 using the fast unfolding of communities in large networks algorithm (version 0.3) and the 100 best (judged by modularity score) were used for downstream analysis. The most informative clustering was selected using gene signature enrichment, visualized using the Gephi package (version 0.82) with the ForceAtlas2 layout method, and an interactive HTML5 web visualisation exported using the sigma.js library (https://github.com/oxfordinternetinstitute/gephi-plugins/tree/sigmaexporter-plugin)(34, 35).

#### Network availability

Interactive networks and meta-data is available at http://pgcna-tgfb.gets-it.net

#### Module overlaps

The overlap of the modules between the networks at the gene and signature level was assessed using a hypergeometric test and the overlap visualized as a python matplotlib heatmap of −log_10_ *p*-values. The signatures were pre-filtered to *p*-value <0.001 and ≥ 5 and ≤ 1000 genes.

#### Heatmap visualizations

Visualizations of the clustered gene expression data and enriched gene signatures were carried out using the Broad GENE-E package (https://software.broadinstitute.org/GENE-E/) data were hierarchically clustered (Pearson correlations and average linkage). For the visualizations of gene signature enrichments, the signatures were filtered to FDR <0.1 and ≥ 5 and ≤ 1500 genes for the signature sets: selecting the top 30 most significant signatures per module; excluding signature sets: MSigDB_C7 and GeneSigDB.

### Ethical approval

Approval for this study was provided by UK National Research Ethics Service via the Leeds East Research Ethics Committee, approval reference: 07/Q1206/47.

## Results

### TGFB supports in vitro PC survival

PC survival can be supported in vitro in a contact-independent fashion by stromal cell lines (6). Amongst the secreted factors released by such cell lines are SDF1 and members of the TGFB family, and in the case of the M2-10B4 cell line, which efficiently supports PC survival in our model system, this is TGFB3 (Supplemental Table 1). Indeed, amongst a series of recombinant factors tested, TGFB3 acted in conjunction with IL6 to support PC survival in vitro, independent of stromal support (Figure 1A and B). Upon TGFB3 stimulation, PB ASCs at day 9 of culture induced robust SMAD2 phosphorylation, which was sustained for at least 4 hours (Figure 1C). This SMAD2 phosphorylation could be inhibited by pretreatment with a TGFB receptor inhibitor consistent with canonical signaling (Figure 1D). Immunohistochemical staining of bone marrow sections identified evidence of nuclear phospho-SMAD2 in PCs (Figure 1E) (36), which was corroborated by flow cytometric analysis of ex vivo bone marrow PCs (Figure 1F and SF1A). Thus, phosphorylation of SMAD2 can be observed as a feature of bone marrow PCs and TGFB provides a potential niche signal capable of driving SMAD2 phosphorylation in primary human ASCs in vitro.

**Figure 1.**
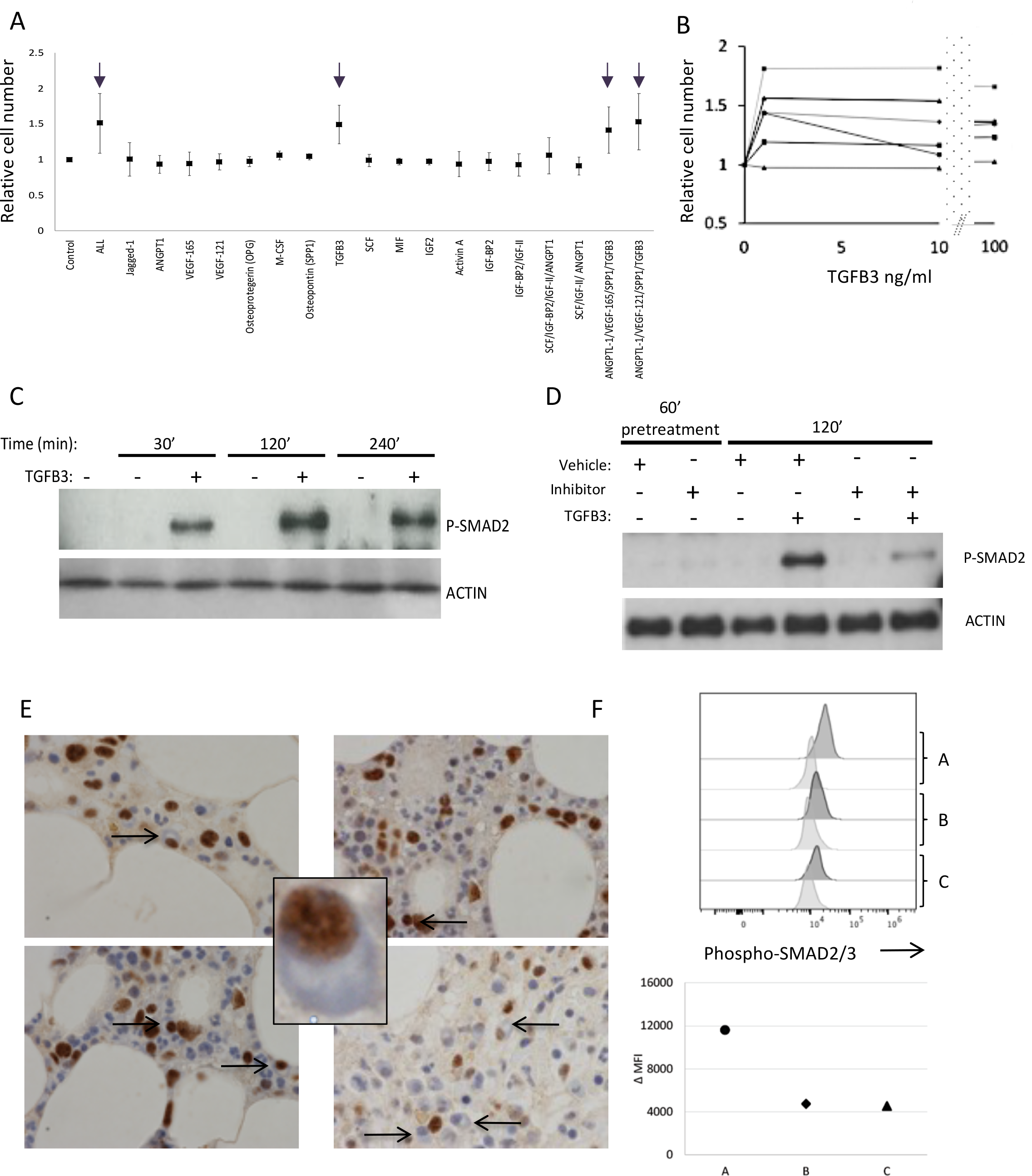
TGFB3 provides a potential survival signal for ASCs. **(A)** A series of proteins, identified from the secretome of stromal cells, was tested for the capacity to promote plasma cell survival in combination with IL6 and IL21. Data are shown as average fold cell number relative to IL6 and IL21 alone (control) from a total of 4 donors in two independent experiments. Proteins tested are indicated on the x-axis, arrows identify all conditions in which TGFB3 was present. **(B)** Dose response data of TGFB3 in ng/ml (x-axis) showing viable plasma cell number at day 13 of culture (y-axis) for five donors. **(C)** Time course of SMAD2 phosphorylation induced by TGFB3 stimulation (2 ng/ml) at day 6 relative to control, representative time points indicated above the Western blot. **(D)** Inhibition of SMAD2 phosphorylation with inhibitor SB525334. Shown is induced SMAD2 phosphorylation at 120min in ASCs in the presence or absence of TGFB3, either pre-treated or not pre-treated with inhibitor for 60 minutes. **(E)** Immunohistochemical detection of phosphorylated SMAD2 in normal human bone marrow trephine samples. Four representative fields are shown with morphologically defined plasma cells identified with black arrows. Insert shows a cropped field to illustrate morphology of a phospho-SMAD2/3 positive plasma cell. **(F)** Intracellular phospho-SMAD2/3 staining in plasma cells gated on CD19 expressing plasma cells showing representative example and ΔMFIfrom three independent bone marrow samples. Isotype control pale grey, phospho-SMAD2/3 dark grey.

## TGFB supports execution of the terminal phase of PC differentiation

The sustained activation of SMAD2 phosphorylation observed following PB stimulation with TGFB3 suggested the potential capacity to impact on subsequent gene expression in PCs. We therefore performed a gene expression time course experiment focusing on the differentiation window of PBs to PCs across sequential time points between day 6 and day 20 of culture in the presence of TGFB3 or IFNα, which represents our previous standard cytokine context (3), or both TGFB3 and IFNα The overall phenotypic maturation of PBs to PCs proceeded without significant differences between the three conditions to day 20 (Figure SF1B), and the numbers of surviving cells at each time point showed no significant difference although exhibiting a trend toward better survival in the presence of IFNα (Figure SF1C).

To analyze the gene expression data (Supplemental Table 2) we first considered differential gene expression comparing each time/condition pair against the day 6 PB state at which the different cytokine conditions were added. Considering genes with fold-change >1.5 and FDR corrected p-value<0.05, we assessed the extent of overlap between sets of repressed and induced genes (Supplemental Table 3) under each condition in all-by-all pairwise comparisons (Figure 2A and C) and as overlapping Venn diagrams (Figure 2B and D). The pairwise comparison heatmaps assess how the sets of significantly downregulated (repressed) or significantly upregulated (induced) genes at a given time point and condition relate to all the sets of significantly upregulated or downregulated genes in either of the other two conditions. This visualization emphasizes the progressive convergence of differentially expressed genes with time. Consistent with our previous analysis (3), a group of differentially up and down regulated genes are characteristic of the response to IFNα (C1 and C2) and are induced within 3h of stimulation regardless of the presence or absence of TGFB3. Thus, the two conditions of differentiation in the presence of IFNα (C1 and C2) are more closely related at earlier time points, particularly for induced genes. Nonetheless, from 24h (day 7) onward there is convergence onto a common set of induced and repressed genes irrespective of the cytokine condition. Venn diagrams assessed the extent of overlap at each time point as a three-way comparison, again illustrating convergence onto a common set of induced and repressed genes over time. However, this also allowed the representation of genes which were unique to each condition, identifying a subset of genes regulated in TGFB3 conditions (Figure 2C and D).

**Figure 2.**
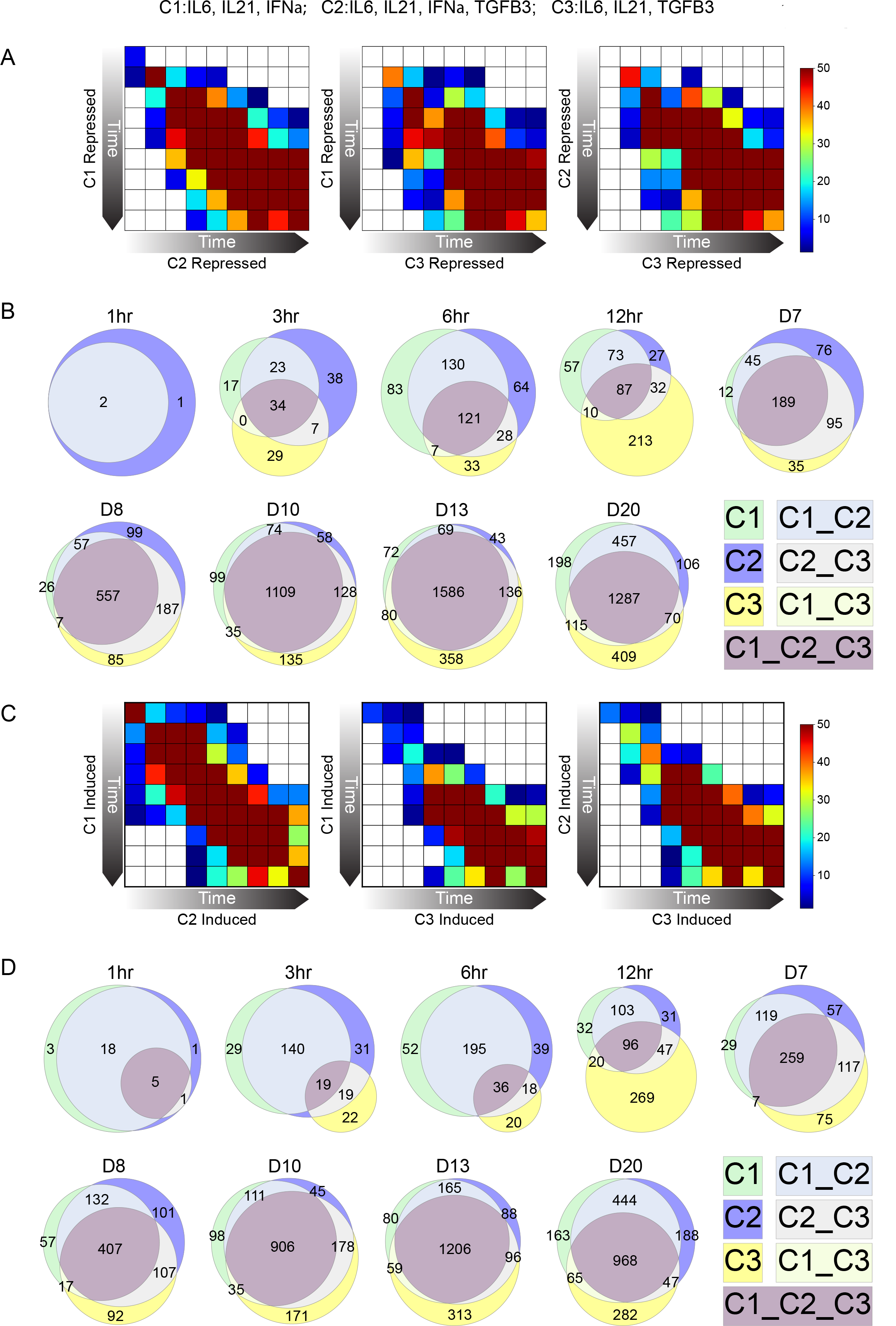
Maturation of PBs to PCs is associated with convergent patterns of gene induction and repression. Genes differentially expressed in each of the three conditions (Cl-IFNα, C2-IFNα + TGFB3, C3-TGFB3 all with IL6 and IL21) at each time point of the time course was determined relative to the initial gene expression at the PB state. The overlap of differentially expressed genes at each time point was then assessed in pairwise fashion and displayed as a heatmap, **(A)** repressed and **(C)** induced, with the significance of overlap as −loglO p-values displayed as a color scale as indicated to the upper right of the figure (white not significant, dark blue p-value < 0.05, to brown highly significant overlap). By comparing the overlap of any given time point to all others in the series the display highlights the overall coordinated progression of differential expression between the conditions. The conditions are indicated on the x- and y-axis labels for each comparison left panel (Cl y-axis vs C2 x-axis), middle panel (Cl y-axis vs C3 x-axis), right panel (C2 y-axis vs C3 x-axis). **(B)** and **(D)** represent the overlaps of differentially repressed and induced genes respectively at each time point, with each condition and intersect color coded as indicated in the figure. The size of the three-way intersect illustrates the progression toward a common set of regulated genes relative to genes uniquely regulated in each condition.

Thus, the time course supports a pattern of convergent gene regulation during the PB to PC transition, whereby irrespective of the culture conditions allowing PC survival a common set of induced and repressed genes emerges. Overlaid onto this are groups of differentially expressed genes which are specific to the stimuli that are permissive to survival and maturation.

## PGCNA resolves discrete biological modules of gene expression

To explore the biology associated with differential gene expression during the PB to PC transition we applied an expression networking approach, which we refer to as parsimonious gene correlation network analysis (PGCNA) (Figure 3A). In parallel work we extensively validate the utility of this method for evaluation of primary cancer expression data sets (33). Here we test its utility in analyzing time course expression data.

**Figure 3.**
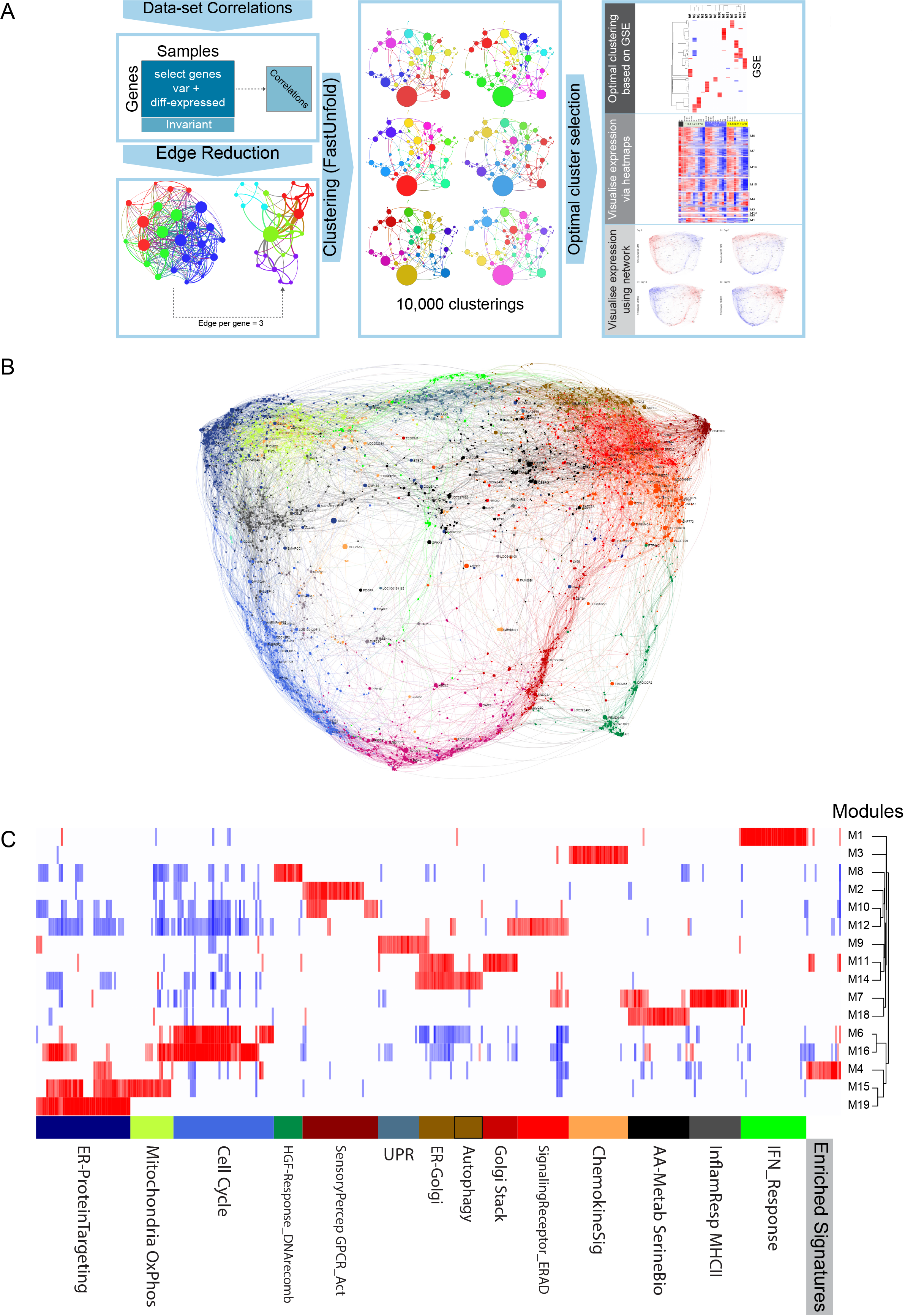
Application of PGCNA to time course gene expression data of PB to PC transition. **(A)** Outline of PGCNA approach as detailed in Methods and (33). **(B)** Network representation of the modular pattern of gene expression during the transition of PB to PC. Network modules are colour coded, for interactive version go to http://pgcna-tgfb.gets-it.net/. **(C)** Heatmap summary representation of gene ontology and signature separation between network modules (filtered FDR <0.1 and ≥ 5 and ≤ 1500 genes; selecting the top 30 most significant signatures per module). Significant enrichment or depletion illustrated on red/blue scale, x-axis (signatures) and y-axis (modules). Hierarchical clustering according to gene signature enrichment. Indicative module terms show below. For high-resolution version and extended data see SF2 and Supplemental Table 5.

The resulting expression network for the transition of PBs into PCs divides into 19 modules (Figure 3B, Supplemental Table 4 and online resource). The separation of specific functions and ontologies between network modules was assessed using enrichment of signature and ontology terms across each module of the network (Figure 3C, SF3, and Supplemental Table 5). The summary display of signature and ontology term enrichments illustrates the resolution of biological processes between modules as discrete bars of enrichment (red) and depletion (blue). These modules in turn reflect differential regulation of these processes across the time course of differentiation (Figure 4). The initial step of the differentiation of PBs into quiescent PCs is accompanied within 3h by an IFN response module (M1_JFNResponse), where this cytokine is present, and this remains in part superimposed on the overall network of gene expression of the differentiating PC at all time points in conditions containing IFNα (Figure 4B and SF4) (3). TGFB3 effects are more distributed within the network including genes in a module linked to both metabolic and translational processes as well as genes linked to the B cell state and the chemokine receptor gene *CXCR4* (M15_RNAProcessing Translation_Mitochondrion_OxPhos).

**Figure 4.**
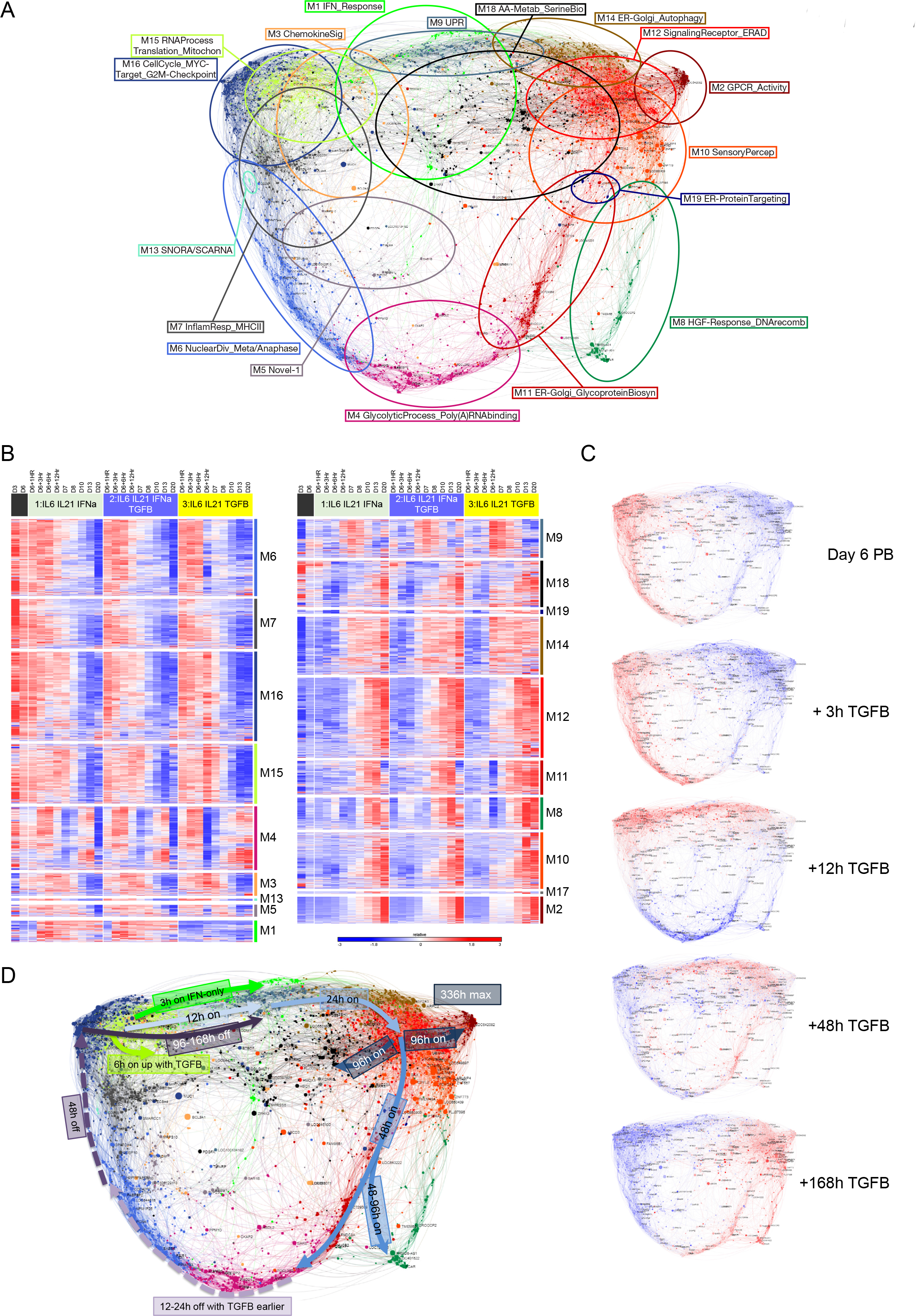
Dynamics of gene expression during PB to PC transition. **(A)** Module summary terms identify different aspects of cell biology represented across the expression network. **(B)** Heat map displaying the pattern of gene expression across the time course for the three conditions (black day 6 reference; light green IL6, IL21, IFNα; blue IL6, IL21, IFNα, TGFB3; yellow IL6, IL21, TGFB3) module numbers indicated on the right, z-score gene expression blue (−3 low) red (+3 high) color scale as indicated right lower edge. Showing the median expression across 4 donors per timepoint, except day 20 where QC failure reduced donor numbers. **(C)** Overlay of gene expression z-scores for all genes in the network shown in blue (low) to red (high) z-score color scale. Day 6 provides the starting reference point for the sequential expression patterns observed under C3:IL6/IL21/TGFB3 condition at the time points indicated to the right of each network image. For interactive versions of all networks go to http://pgcna-tgfb.gets-it.net/. **(D)** Summary of gene expression flow across the time course illustrated with arrows in the network. Induced expression in solid lines, repression in dotted lines, with relevant time points and conditions indicated in the figure.

Following the initial response to cytokine conditions, a module linked to genes regulated by IRE1 and the unfolded protein response and overlapping with ER/Golgi components is upregulated (M9_UPR) (Figure 4B and C and SF4). This initial ER response is relatively accelerated in the presence of TGFB3 with greatest expression at 12h rather than 24h. In all conditions, this is followed by sequential modules of gene expression which separate secretory pathway components and genes linked with autophagy (M14_ERGolgi_Autophagy) from those enriched for ERAD and ER quality control pathways (M12_SignalingReceptor_ERAD) and those linked to the Golgi apparatus and glycoprotein biosynthesis (M11_ER-Golgi_GlycoproteinBiosyn). Thus, as ASCs progress from PB to the PC state, modules of genes indicative of an ER stress response precede those linked to optimization of secretory activity.

The induction of ER-related modules is accompanied on the other hand by progressive gene silencing. This initiates with the repression of genes linked to a subset of cell cycle related processes, in particular those related to the mitotic anaphase and sister chromatid segregation at 12-24h (M6_NuclearDivision_Meta/Anaphase). Other components of the cell cycle remain expressed up to 24h after transition into PC supportive conditions (M16_CellCycle_MYCTarget_G2MCheckpoint). The initial extinction of cell cycle-related components is modestly accelerated in the context of TGFB3 (Figure 4B (M6 & M16) and Figure SF4).

Subsequently, from 48h onward the main cell cycle and DNA replication module (M16_ CellCycle_MYCTarget_G2MCheckpoint) and a module linked to antigen presentation via MHC class-II and inflammatory signaling (M7_InflammatoryResponse_MHCII) are repressed. This is followed by the eventual repression of the UPR gene module (M9_UPR), initially induced at the start of the transition to the PC state.

The late phase, beyond 96h (day 10), is accompanied by the progressive concentration of PC-related gene expression into secretory and alternate metabolic related modules including those linked to amino acid metabolism (M18_AminoAcidMetab_SerineBiosyn). Distinctively mature PC-associated modules are enriched for microRNA and non-coding RNA genes as well as subsets of signaling receptors including G-protein coupled receptors (M2_GPCR_Activity).

This network analysis of the PB to PC transition provides a refined view of the final steps of the differentiation of ASCs, linking gene expression related to secretory optimization to cell cycle exit and expression of select surface receptors. Exposure to TGFB3, unlike IFNα, did not impose a distinct single module of gene expression in the conditions necessary for this long-term time course, instead exerting more distributed effects across several modules.

## TGFB modulates SDF1 signaling to the ERK MAP kinase pathway

Amongst the genes differentially expressed in response to TGFB3 at the early phase of the response and subsequently sustained was *CXCR4* (Figure 5A), which is a known target of TGFB signaling in other cellular contexts (16, 17). Since SDF1-CXCR4 signaling is considered to be a primary determinant of recruitment and residence of PCs in the bone marrow niche (11–13), we focused further on this aspect of the response.

**Figure 5.**
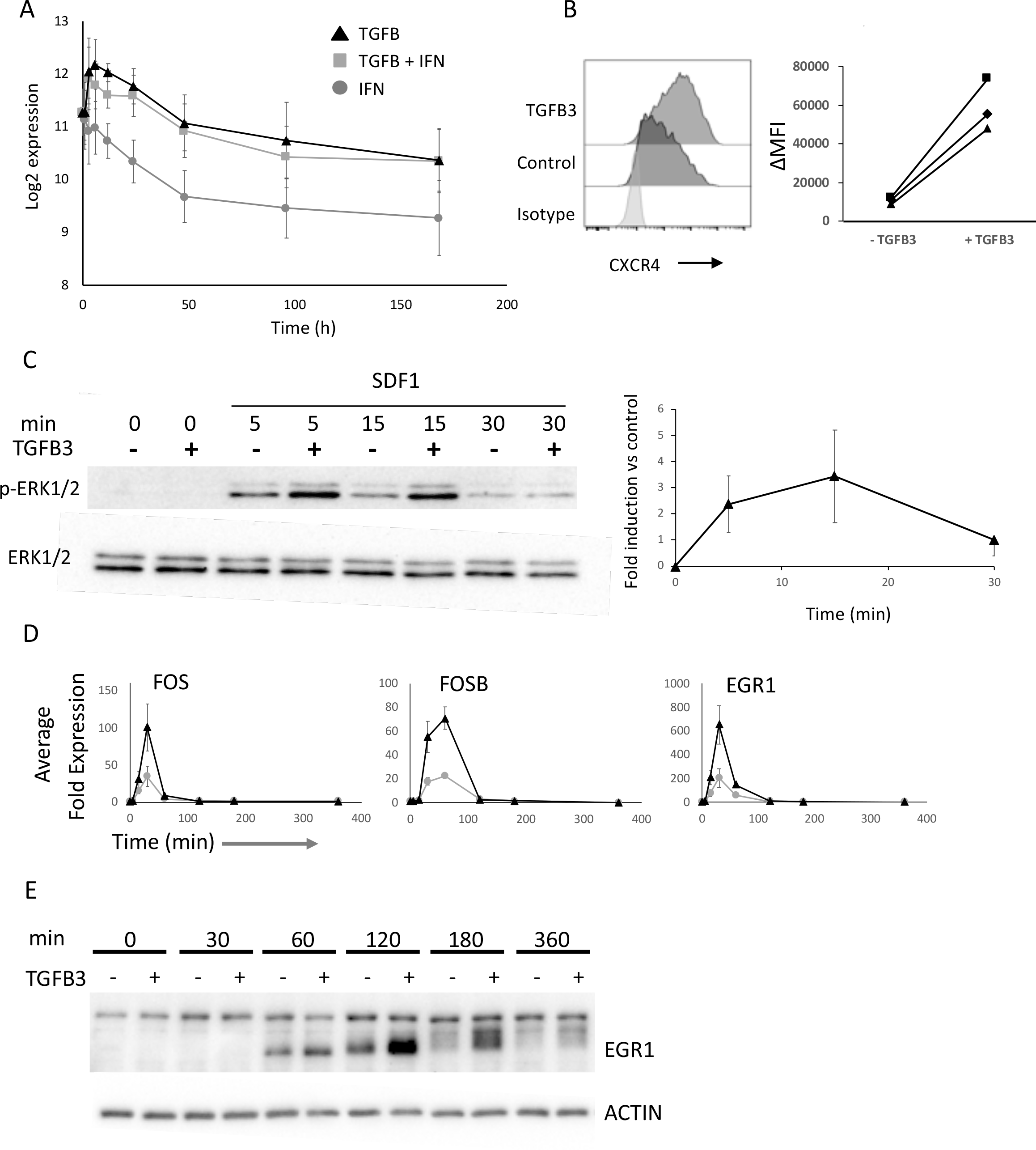
TGFB3 induces CXCR4 expression in ASCs. **(A)** Expression of *CXCR4* mRNA across B-cell differentiation showing average and standard deviation (n=4 samples per time point and condition) of log-2 gene expression values for the three conditions TGFB3(black triangle), TGFB3 + IFNα (grey square), IFNα (grey circle) each in the presence of IL6 and IL21. **(B)** Representative data for CXCR4 surface expression on ASCs at day 7 of culture in the presence (mid grey upper contour) or absence (dark grey middle contour) of TGFB3 relative to isotype control light grey (lower contour) in control treated samples (light grey). Summary of change in CXCR4 MFIfor three representative donors. **(C)** Time course of ERK phosphorylation induced by SDF1 treatment from 0-30min in the presence (+) or absence (−) of prior TGFB3 exposure as indicated above the blot, upper panel Western blot for phospho-ERKl/2, lower panel Western blot for total ERK1/2. Quantitation of fold ERK1/2 phosphorylation across the time course of TGFB3 treated vs untreated control samples (3 independent replicates). **(D)** Quantitation of *FOS*, *FOSB* and *EGR1* mRNA expression following SDF1 treatment of ASCs in the presence (black triangle) or absence (grey circle) of prior TGFB treatment, average and standard error of mean from three replicates. **(E)** EGR1 protein expression across the indicated time course following SDF1 treatment in ASCs in the presence (+) or absence (−) of prior TGFB3 treatment as indicated. Lower panel ACTIN loading control.

CXCR4 is expressed on the surface of PBs and PCs and was increased on PBs following treatment with TGFB3 (Figure 5B). SDF1 signaling via CXCR4 can mediate a variety of intracellular signaling events. Amongst these is activation of the MAP kinase pathway (21, 37). Indeed, treatment of ASCs with SDF1 at day 7 of culture led to rapid induction of ERK1/2 phosphorylation (Figure 5C). Furthermore ERK1/2 phosphorylation induced in response to SDF1 was amplified and sustained by pre-treatment with TGFB3 (Figure 5C). Receptor density has been identified as a means by which the MAP kinase pathway output from growth factor receptors can be modified (27, 29), and these data would be consistent with such a model.

To evaluate whether the enhanced ERK1/2 activation impacted on gene regulation we examined the expression of immediate early genes *EGR1*, *FOS* and *FOSB.* Following SDF1 stimulation, PBs exhibited a rapid induction of *FOS* and *EGR1* at 15 and 30 min with decay by 60 min and return to near baseline by 120 min (Figure 5D). *FOSB* showed a slightly delayed kinetics with initial upregulation at 30 min and peak at 60 min. For each of these genes pre-treatment with TGFB3 substantially increased the amplitude of the response in all donors, albeit with variability of the absolute values between donors. Thus, the primed state of such immediate early response genes is retained in PBs as at other stages of the B cell lineage (38).

EGR1 is a transcriptional regulator implicated in PC biology through its recurrent mutation in PC myeloma (23–25). Furthermore, EGR1 provides a potential sensor of the duration of MAP kinase signaling, whereby persistence of ERK1/2 activation allows phosphorylation of EGR1, stabilizing its expression and impacting on downstream function (27). We therefore examined the expression of EGR1 at protein level following SDF1 treatment in the presence or absence of prior TGFB3 exposure. SDF1 treatment, without prior TGFB3 exposure, induced expression of EGR1 protein which was detectable by 60min, peaked at 120min, and returned to near baseline at 180min. By contrast, SDF1 treatment following prior TGFB3 exposure substantially increased the magnitude of EGR1 protein expression at both 120 and 180min. EGR1 protein expression remained significantly above baseline at 360min (Figure 5E). This impact on EGR1 expression was linked to an altered mobility in the EGR1 band, which is consistent with the effects of phosphorylation mediated by persistent ERK1/2 activation (27).

Thus, EGR1 is dynamically regulated in response to SDF1 in primary human ASCs. The pattern of regulation is consistent with a mechanism whereby enhanced and sustained ERK signaling is linked both to the regulation of the magnitude of gene expression and to subsequent protein phosphorylation and stabilization (28, 29).

## SDF1 induces a growth factor-like gene regulatory response in ASCs

To provide a more global picture of the impact of SDF1 signaling in ASCs in the presence or absence of TGFB3 we again utilized a gene expression time course approach. Differentiating PBs were maintained in low serum media, with or without TGFB3 for 20h prior to acute stimulation with SDF1. The response was sampled at baseline immediately prior to SDF1 exposure and 30, 120 and 360min after stimulation to catch the dynamics of immediately early and delayed response gene expression (Supplemental Table 6) (39).

For this analysis we initially considered differentially expressed genes with FDR corrected p-value <0.05 (Supplemental Table 7). After 20h TGFB3 treatment 12 genes were significantly upregulated and 5 genes significantly downregulated relative to the control treated sample. Amongst these was *CXCR4*, which remained significantly differentially expressed at all times point of the experiment. Following addition of SDF1 a pulse of differentially expressed genes was observed, which was broader in the presence (33 genes significantly induced) than in the absence of TGFB3 (14 genes significantly induced) relative to baseline for the respective condition at 30min. All but one of the 14 genes significantly induced following SDF1 treatment in the absence of TGFB3 were encompassed in the 33 genes induced in the presence of TGFB3, this included *EGR1*, *FOS* and *FOSB* as well as *ATF3*, *EGR2*, *CD69* and *KLF6.* In the presence of TGFB3 additional genes significantly upregulated included *JUNB*, *M1R155HG* and *SRF.* At 120 min after SDF1 treatment in the presence or absence of TGFB3, 154 and 44 genes were significantly upregulated, and 172 and 18 genes significantly downregulated, respectively. Again, the majority of the genes regulated in the absence of TGFB3 were included amongst genes regulated in the presence of TGFB3, consistent with a substantially diversified response in cells exposed to TGFB3. In both conditions, by 360min after SDF1 stimulation the response was curtailed.

We next considered the wider pattern of gene expression change induced in this model using PGCNA and a lenient threshold for differential expression. This resulted in a network comprised of 16 modules (Figure 6A, Supplemental Table 8, and online resource). Gene ontology and signature enrichment analysis indicated that these modules were associated with coherent biology suggesting that additional insight into the overall response could be derived by integrating subtle changes across many genes in this manner (Figure 6B, SF6, and Supplemental Table 9).

**Figure 6.**
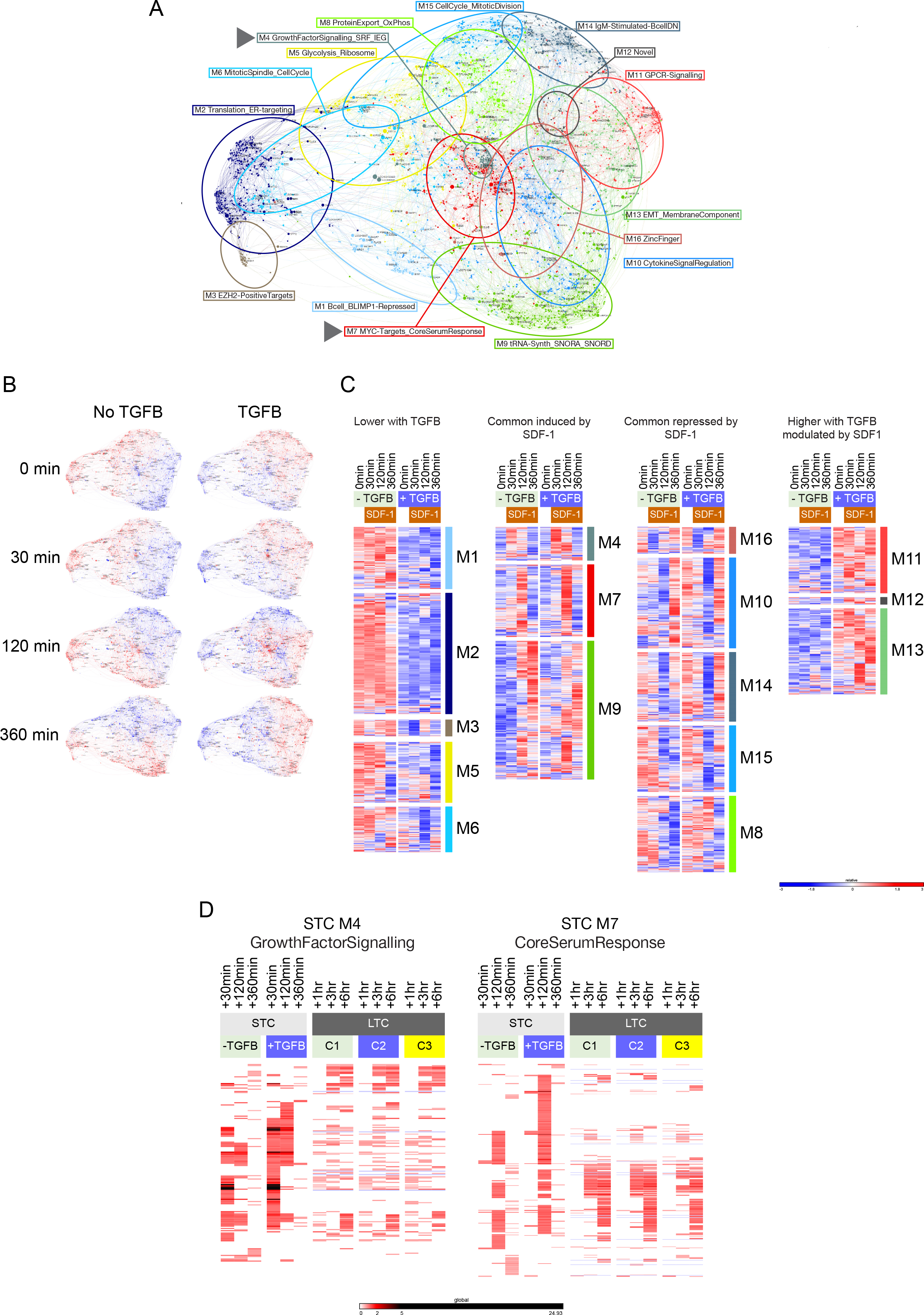
Network analysis of SDF1 induced gene expression in ASCs. **(A)** PGCNA analysis of gene expression derived from a short time course (STC) following SDF1 treatment in presence or absence of TGFB3. Network comprises 16 modules, signature enrichment analysis was used to determine biological features of individual modules and derive representative summary terms. Arrow heads identify modules M4 (GrowthFactorSignalling) and M7 (CoreSerumResponse). **(B)** Overlay of gene expression z-scores for genes in the network shown in blue (low) to red (high) z-score color scale. Conditions are indicated above the network images with time point of sampling in relation to SDF1 treatment to the side (for interactive networks see go to http://pgcna-tgfb.gets-it.net/; detailed version SF5). **(C)** Module expression values illustrated as heat map separated into categories of lower with TGFB, induced following SDF1, repressed following SDF1 and higher with TGFB. Showing the median expression across 3 donors per timepoint. **(D)** Heat map illustrating dynamics of M4 and M7 gene expression derived from the short time course (STC) and comparing patterns in the short time course data to the long time course (LTC) data with the relevant conditions C1, C2 and C3 indicated in the figure. Shows expression fold change for each gene relative to baseline (STC: 0min/day7, LTC: day6).

Prior to SDF1 exposure, modules of co-expressed genes differentiated the baseline states with or without TGFB3. Cells cultured in the absence of TGFB3 showed generally higher expression of several modules including one enriched for BLIMP1 target genes (M1_Bcell_BLIMP1Repressed), genes involved in translation (M2_Translation_ER-Targeting), and genes linked to the mitotic spindle and cell cycle (M6_MitoticSpindle_ CellCycle). By contrast, cells cultured in the presence of TGFB3 for 20h showed relative upregulation of gene modules linked to GPCR signaling (M11_GPCR-Signalling), and integral membrane proteins (M13_EMT_MembraneComponent), including *CXCR4.* Thus, TGFB3 exposure for 20h promoted a subtle shift toward a more differentiated phenotype in the ASC population. Cells under both conditions, prior to SDF1 exposure, retained elements of cell cycle associated gene expression (M15_CellCycle_MitoticDivision) and expression of genes linked to protein export and oxidative phosphorylation (M8_ProteinExport_0xPhos).

Against this backdrop, PGCNA indicated a coordinated response of up- and downregulated gene expression following SDF1 treatment. At 30 min this was characterized by the induced expression of genes focused in a single module enriched for characteristics of growth factor responses, targets of serum response factor, and immediate early genes (M4_GrowthFactorSignalling_SRF_IEG) (Figure 6B and C). A second module of delayed responses genes was induced at the 120min time point, again with greater magnitude in the TGFB3 treated cells, which was enriched for core serum response associated genes (M7_MYC-Targets_CoreSerumResponse). A further module (M9_tRNA-Synth_SNORA_SNORD) was upregulated at 360min, consistent with the kinetics of secondary response genes, and included genes associated with amino acyl tRNA synthesis, SNORA/SNORD genes and the antioxidant genes *HMOX1*, *NQO1* and *GCLM.* The sequence of common upregulated modules was also paralleled in modules of sequential gene repression following SDF1 exposure at 30 and 120min with subsequent re-expression.

The analysis with PGCNA thus emphasized a sequence of gene expression changes following SDF1 exposure with strong parallels to that of growth factor responses, indicating that such a regulatory response can be driven by this niche signal in ASCs. To assess whether induction of a growth factor like signaling module was a unique feature of the SDF1 response we re-examined the long time course data set focusing on the pattern expression of genes belonging to the growth factor response and core serum response modules induced by SDF1 (M4 and M7) (Figure 6D). While a core component of IEG including *FOS*, *FOSB*, *JUN*, *EGR1*, *EGR2* and *EGR3* were acutely induced by signals promoting survival this was to a more modest degree than observed in response to SDF1 and did not extend across the M4 module genes. In contrast, a larger proportion of the M7 module genes showed related patterns of regulation at +3h and beyond. We conclude that a pulse of growth factor-like signaling can be delivered in ASCs, in particular exemplified by the acute response to SDF1, but that this response is not an intrinsic requirement for PC survival, at least as assessed in vitro.

## Discussion

The data presented here support the conclusion that TGFB can act as part of a realized in vitro PC niche and in this context can mediate cross-talk with the SDF1-CXCR4 pathway. The demonstration that TGFB stimulation alters expression of CXCR4 and the nature of signaling responses to SDF1 is shared with other cell types (16,17). Here we have dissected this response in ASCs in detail and identified EGR1 and immediate early genes as a proximal point of signal integration upstream of subsequent waves of gene expression. Furthermore, by comparing the modular patterns of gene expression induced during cultures supporting PC survival and those following acute SDF1 exposure we find that growth factor-like gene regulation is separable from sustained gene expression associated with ASC survival/maturation.

The data additionally provide insight into the sequence of gene expression changes during the final stage of human ASC maturation from the PB to the PC state. To address this question, we have applied a method we have developed, which we refer to as parsimonious gene correlation networking analysis (PGCNA), to gene expression time course data. PGCNA resolves a detailed sequence of expression modules that are both induced and repressed as the ASC completes its maturation to the quiescent PC state. In the context of our long time course data of maturation to the PC state this illustrates a wave of UPR-related gene expression as an initial feature of this differentiation step. This is eventually extinguished but precedes the upregulation of several distinct modules of secretory pathway-related gene expression. This is consistent with a model in which the optimal adaptation for secretory capacity is not completed at the phenotypically defined PB ASC stage, and that a classical UPR response accompanies the final stages of differentiation to the quiescent PC state. Indeed, in murine PC differentiation the UPR transcription factor XBP1 is largely dispensable for the earlier phenotypic maturation but is essential for the optimization of PCs for maximal secretory activity (40).

In relation to TGFB signaling our data illustrate that classical SMAD phosphorylation is readily activated and sustained in ASCs, but that the overall effects on gene expression are modest. Low serum presents a limiting factor for long-term PC cultures but in short-term cultures this is not the case, and in this context analysis of subtle gene expression changes with PGCNA indicates that TGFB3 treatment accelerated extinction of BLIMP1 target genes, and elements of the cell cycle. These features are consistent with the established role of TGFB as a factor capable of contributing to the establishment of cell cycle quiescence in other systems (41).

The SDF1-CXCR4 axis represents one of the main determinants of PC niche homing (11–13). Here we show that this niche signal in human ASCs is coupled to activation of the ERK MAP kinase pathway impacting on immediate early gene expression. Indeed, SDF1 exposure produces a pattern of gene expression closely related to that of classical growth factor signaling (39). By contrast, in the long term culture a distinct immediate early gene/growth factor-like signaling module is not identified. Nevertheless, a core subset of immediate early genes is acutely induced but not maintained after initial exposure to conditions promoting plasma cell maturation and survival. Thus, in ASCs the induction of a pulse of growth factor-like signaling is induced by niche signals related to homing and survival but is rapidly attenuated and separated from gene expression associated with survival and maturation. Transient and sustained pulses of growth factor-like signaling distinguish proliferation from differentiation promoting growth factors in models such as the PC12 neuronal cell line (29). Speculatively, the modulation of MAP kinase signal intensity and duration may also play a significant role in PC biology. In general, the importance of MAP kinase signaling in PC biology is suggested by the recurrent mutations of *NRAS*, *KRAS* and *BRAF* in myeloma (23–25). While the differential signaling in ASCs consequent on such mutations remains largely unexplored, a general prediction is that these impact on the kinetics and amplitude of MAP kinase pathway activation. Mutations affecting this pathway also extend to recurrent mutations in *EGR1.* Indeed, in a large recent analysis it was shown that when EGR1 mutations occur in myeloma these show a high clonal fraction suggesting that the mutation either exerts a strong selective pressure or occurs as an early clonal event (5). The data presented here indicate that a growth factor-like signaling pathway impacting on EGR1 expression is a feature of the acute response to niche signals but is not necessarily sustained during PC survival. Furthermore, our data indicate that enhanced MAP kinase signaling can act in ASCs to sustain EGR1 protein expression. It will be interesting in future to explore what effects mutations of upstream regulators and EGR1 itself have on EGR1 behavior in PC, and how these may intersect with the growth factor like response induced by niche factors.

In conclusion, using an in vitro model system of primary human ASC differentiation, and applying a gene expression networking approach, the data presented here argue that MAP kinase signaling and immediate early gene regulation can be imposed by niche signals onto the ASC expression profile, but are not sustained as a module integral to PC survival. ASCs can encode acute niche signals through modulation of the intensity and duration of immediate early gene regulation, with EGR1 providing an example of a point of niche signal integration.

## Acknowledgements

This work was supported by Cancer Research UK program grant (C7845/A17723). We thank Ulf Klein for critical review of the manuscript.

## Author contribution

SS, performed experiments, analyzed data, wrote papers, MAC performed analyses, analyzed results, conceived study, wrote paper, IF and AZ performed experiments DRW guided analyses, edited paper, GMD analyzed results, wrote paper, RMT conceived study, analyzed results, wrote paper.

## Supplemental Figure Legends

**SF1. Accompanies Figure 1 and 2**. **(A)** Gating strategy for identification of bone marrow PCs based on FSC, SSC, CD38, CD19, CD138 and CD56 to separate CD19+ PCs from B cells and CD56+ expressing potential neoplastic plasma cells. **(B)** Phenotypic maturation of ASCs in the long-term culture under the three conditions used for gene expression analysis. Figure shows plots of flow cytometry data for CD19 vs CD20 (left panel of each pair) and CD38 vs CD138 (right panel of each pair). At day 6 (top left) and then for each of the three conditions tested at day 7, 8,10,13 and 20 as indicated beneath the flow plots. The conditions are identified on the left of the figure. **(C)** Viable cell numbers derived from 7AAD staining and bead-based quantification at each time point of the culture system shown as average and standard deviation of three donors. IL6, IL21, IFNα (grey circle and line), IL6, IL21, IFNα, TGFB3 (open square dotted line), IL6, IL21, TGFB3 (black triangle, solid line).

**SF2. Accompanies Figure 3**. High resolution version of Figure 3C. Heatmap of gene ontology and signature term enrichments linked to the PGCNA modules of the time course network analysis (filtered FDR <0.1 and ≥ 5 and ≤ 1500 genes; selecting the top 30 most significant signatures per module). For full signature enrichment lists, please see Supplemental Table 5. Modules are shown along the x-axis, and selected signature terms along the y-axis. Signature terms and modules are hierarchically clustered to illustrate relationships. Enrichment (red) and depletion (blue) of signatures are shown on color scale of z-score.

**SF3. Accompanies Figure 4**. Network representation of dynamics of gene expression during PB to plasma cell transition in the presence or absence of TGFB3. Overlay of gene expression z-scores for all genes in the network shown in blue (low) to red (high) color scale. The expression state at day 6 provides the common reference with the expression patterns at each time point for each of the three conditions shown to the right. IFNα, IFNα and TGFB3, or TGFB3, each in the context of IL6 and IL21, shown as indicated. The time point information is illustrated across the figure. Interactive versions of all networks are available on line.

**SF4. Accompanies Figure 6**. Heatmap of gene ontology and signature term enrichments linked to the PGCNA modules of the SDF1 time course network analysis (filtered FDR <0.1 and ≥ 5 and ≤ 1500 genes; selecting the top 30 most significant signatures per module). For full signature enrichment lists, please see Supplemental Table 9. Modules are shown along the x-axis, and selected signature terms along the right-hand side y-axis. Signature terms and modules are hierarchically clustered to illustrate relationships. Enrichment and depletion of signatures are shown on a red to blue scale.

**SF5. Accompanies Figure 6**. **(A)** Network illustration with modules identified by respective summary terms. **(B)** Representation of dynamics of gene expression during SDF1 response in presence or absence of TGFB3. Gene expression z-scores for all genes in the network are overlaid on network elements as blue (low) to red (high) color scale. Time points are shown on the left, with conditions no TGFB left, TGFB right. Annotated network with module summary terms is provided above.

